# Deletion of KLHDC3, an E3 ubiquitin ligase complex substrate receptor, leads to obesity in mice

**DOI:** 10.1101/2025.08.05.668783

**Authors:** Paula Armina V. Buco, Ashfaqul Hoque, Wilson Castillo-Tandazo, Alistair M. Chalk, Monique F. Smeets, Carl R. Walkley

**Affiliations:** Centre for Innate Immunity and Infection Diseases, Hudson Institute of Medical Research, Clayton, Victoria 3168, Australia; St. Vincent’s Institute of Medical Research, Fitzroy, Victoria 3065, Australia; Department of Medicine, Melbourne Medical School, University of Melbourne, Fitzroy, Victoria 3065, Australia; Department of Molecular and Translational Sciences, Monash University, Clayton, Victoria 3168, Australia; Drug Discovery Biology, Monash Institute of Pharmaceutical Sciences, Faculty of Pharmacy and Pharmaceutical Sciences, Monash University, Parkville, Victoria 3052, Australia

**Keywords:** KLHDC3, C-end degron, DesCEND pathway, ubiquitination, obesity

## Abstract

Protein ubiquitination is a critical post-translational modification that regulates protein stability and cellular homeostasis. KLHDC3 is a substrate recognition receptor within the recently identified C-terminal degron-mediated DesCEND ubiquitination pathway. It has been characterised to selectively bind proteins with C-terminal RxxxG motifs, targeting them for degradation. Unlike the well-characterised N-terminal degron pathway, the physiological roles of C-terminal degrons remain poorly understood. To investigate KLHDC3 function *in vivo*, we generated Klhdc3-deficient (*Klhdc3^-/-^*) mice. These mice exhibited sub-Mendelian birth rates and progressive postnatal lethality, with a median survival of 136 days and maximum survival observed of ∼1 year of age. Early growth retardation was apparent, followed by a normalisation of body mass with age. However, as *Klhdc3^-/-^* mice aged they developed pronounced obesity at the expense of lean mass, with some individuals reaching fat mass exceeding 50% of total body weight. Combined transcriptomic and proteomic analyses of *Klhdc3^-/-^* embryonic fibroblasts revealed significant changes in protein expression with minimal impact on transcript levels, consistent with KLHDC3’s role in post-translational regulation. Among the upregulated proteins, HINT1 was identified as a novel KLHDC3 substrate, possessing a C-terminal degron motif. Protein stability assays and immunoblotting confirmed HINT1 as a direct target of KLHDC3. These findings establish a new *in vivo* physiological role for the DesCEND pathway and highlight KLHDC3 as a key regulator of development, survival, and adiposity in mice.

## Introduction

The ubiquitin system regulates the post-translational modification of proteins by ubiquitin or ubiquitin–like proteins, and is essential for cellular and organismal homeostasis (Hershko & Ciechanover, 1998). The ubiquitination of proteins can regulate stability, activity either positively or negatively, and sub-cellular localisation of the targeted substrate. This process plays a vital role in various cellular processes including protein homeostasis, DNA repair and cell signalling. Deregulation of proteostasis and protein degradation is associated with a range of disorders, ranging from cancers and immune phenotypes through to neurodegenerative disease. This pathway is established as a therapeutic target, particularly in cancer (Manasanch & Orlowski, 2017), through small molecule inhibition of the proteasome and more recently via the rapidly emerging field of targeted protein degradation (TPD) through PROTAC (proteolysis targeting chimera) molecules (Bekes *et al*, 2022; Nalawansha & Crews, 2020; Zhou *et al*, 2025).

The most common form of ubiquitination occurs through the sequential activity of three enzymes: firstly, ubiquitin is activated by an E1 enzyme, then transferred to an E2 ubiquitin-conjugating enzyme, and finally attached to the target substrate that has been recruited to the complex by an E3 ubiquitin ligase (Timms & Koren, 2020). The specificity of the ubiquitin-proteasome system is achieved via a large number of E3 ligases. E3 ligases can have varied functions, acting in proteostasis as part of quality control of the proteome, while others function in regulating signal transduction cascades (Cruz Walma *et al*, 2022; Zheng & Shabek, 2017). E3 ligases of the ubiquitin-proteasome pathway interact with protein substrates via recognition of distinct molecular sequences/motifs, termed “degrons”. These degron motifs can be present in native proteins or arise from errors during protein synthesis, or become exposed due to protein misfolding or when protein complexes do not form correctly (Timms & Koren, 2020). The first identified and best-characterised degrons are found at the N-termini of the proteins, termed the “N-degron” or “N-end rule” (Bachmair *et al*, 1986; Bartel *et al*, 1990; Sherpa *et al*, 2022). In addition to the prevalent “N-end rule” ubiquitin pathway, internal degron and more recently, C-terminal degron pathways termed destruction via C-end degrons (DesCEND), have been described (Koren *et al*, 2018; Lin *et al*, 2015; Lin *et al*, 2018; Sherpa *et al*., 2022).

The DesCEND pathway is affected by a diverse set of E3 ligases. The C-end degron targeting E3 ligases all utilise tandem repeat domains to recognise the specific motif required for degradation. The main families of DesCEND E3 ligases are the Kelch repeats, Ankyrin repeats, WD repeats, Armadillo-like repeats, and TPR (tetratricopeptide) repeats (Timms & Koren, 2020). The E3 ligases engage Elongin-B, Elongin-C and Cullin 2 (CUL2) (for Kelch, TPR and Ankyrin repeats) or utilise DNA damage-binding protein 1 (DDB1) and CUL4 (for WD and Armadillo-like repeats) (Timms & Koren, 2020). Since the identification of this pathway, a range of substrates have been identified that have specific C-end degrons that mediate targeting by specific E3 ligases including both epitopes in native proteins and aberrant protein products (Koren *et al*., 2018; Lin *et al*., 2018; Makaros *et al*, 2023; Rusnac *et al*, 2018; Scott *et al*, 2024a; Scott *et al*, 2024b; Scott *et al*, 2023; Timms *et al*, 2023; Timms *et al*, 2019; Zhang *et al*, 2023). There is a growing appreciation of the potential of the DesCEND pathway, however, our understanding of the physiological functions and *in vivo* functions of this pathway remains limited.

The KLHDC family of E3 ligases contains 10 members (Gupta & Beggs, 2014; Pilcher *et al*, 2025). The members have been characterised to varying extents, with KLHDC2 being the most extensively described (Rusnac *et al*., 2018; Scott *et al*., 2024a; Scott *et al*., 2024b; Scott *et al*., 2023). There is growing interest in the application of KLHDC members, particularly KLHDC2, for use in targeted protein degradation (Hickey *et al*, 2024; Kim *et al*, 2023; Roth *et al*, 2023; Zhou *et al*, 2024). Understanding the essential physiological functions and identifying substrates for each specific KLHDC member will be important for exploiting these E3 ligases in future. However, predicting substrates based on the degron motifs has so far proven ineffective, due to the relatively loose consensus sequence (Yeh *et al*, 2021). Additionally, emerging data suggest that cryptic KLHDC targeted peptides may be hidden within the primary protein sequence (Yeh *et al*., 2021). Recent large-scale efforts have begun to provide greater understanding of the degron motifs for many KLHDC members (Scott *et al*., 2023; Timms *et al*., 2023; Zhang *et al*., 2023), but we do not have *in vivo* models for the majority to understand these functions in the context of a whole organism. Understanding the essential physiological functions and identifying substrates for each specific KLHDC member will be important for exploiting these E3 ligases in future.

We became specifically interested in KLHDC3 (also known as Peas or 1300011D16Rik) after identifying it in a forward genetic screen (Buco *et al*, 2025). KLHDC3 was originally identified as a testis-specific RAG2-like protein with expression in spermatocytes and localising in the cytoplasm and on meiotic chromatin (Ohinata *et al*, 2003a). Using cell line-based global protein stability (GPS) reporter assays (Huang *et al*, 2023), it was established that KLHDC3 preferentially binds to substrates with C-end RxxxG motifs, promoting their ubiquitination and degradation (Timms *et al*., 2023; Yeh *et al*., 2021; Zhang *et al*., 2023). The structural basis for the C-degron selectivity of KLHDC3 was recently reported (Scott *et al*., 2024a), however, the physiologically relevant substrates of KLHDC3 remain largely unknown, except for p14^ARF^ and c-Myc in human cancer cell lines (Liu *et al*, 2022; Motomura *et al*, 2024; Yeh *et al*., 2021; Zhang *et al*, 2022). KLHDC3 may be important in the lifecycle of bovine herpesvirus, indicating a role in pathological processes that are not at present well understood (Slusarz & Lipinska, 2024; Wachalska *et al*, 2024). New approaches for the identification of substrates are required and to understand the physiological relevance of the C-degron pathways. To this end, we describe here the generation and analysis of *Klhdc3*-deficient mice. *Klhdc3* loss was investigated in both whole animals and cell line models. The *in vivo* phenotypic characterisation of *Klhdc3*^-/-^ mice, supplemented with transcriptomic and proteomic analysis of *Klhdc3*^-/-^ derived cell lines, revealed a role for KLHDC3 in suppressing adiposity/obesity *in vivo* and identified HINT1 as a new physiological substrate.

## Results

### Generation of a *Klhdc3*-deficient allele in mice

The *Klhdc3* gene in mouse is located on chromosome 17:46985476-46991840 (-strand; Ensembl annotation GRCm39) and is composed of 13 exons, with a predicted protein length of 382 amino acids and a molecular mass of 43,088 Da (Fig 1A, 1C). To generate a KLHDC3-deficient mouse line on the C57BL/6J background we used CRISPR/Cas9 to insert loxP elements flanking *Klhdc3* exons 4 and 5 (Fig 1B-1D) (Buco *et al*., 2025). During this process we identified an incidental mutation arising from a fusion between the sgRNA-induced breaks. This resulted in a 676bp deletion that removed exons 4 and 5 of *Klhdc3* (the same exons being flanked by loxP elements), yielding a constitutive heterozygous deletion of *Klhdc3* (Fig 1B). Correct copy number and genomic modification were confirmed by locus-specific PCR and Sanger sequencing. This male was bred to C57BL/6J females to establish a germ-line *Klhdc3^+/-^* allele. Offspring were then bred to a second generation to confirm germ-line transmission, and a breeding colony was established from inbreeding of the F2 generation. We also identified founders where the loxP elements had been successfully and correctly integrated. This conditional *Klhdc3^fl/fl^* allele containing loxP flanked exon 4 and 5 has been used in a separate study and is not further described here (Buco *et al*., 2025).

**Figure 1.**
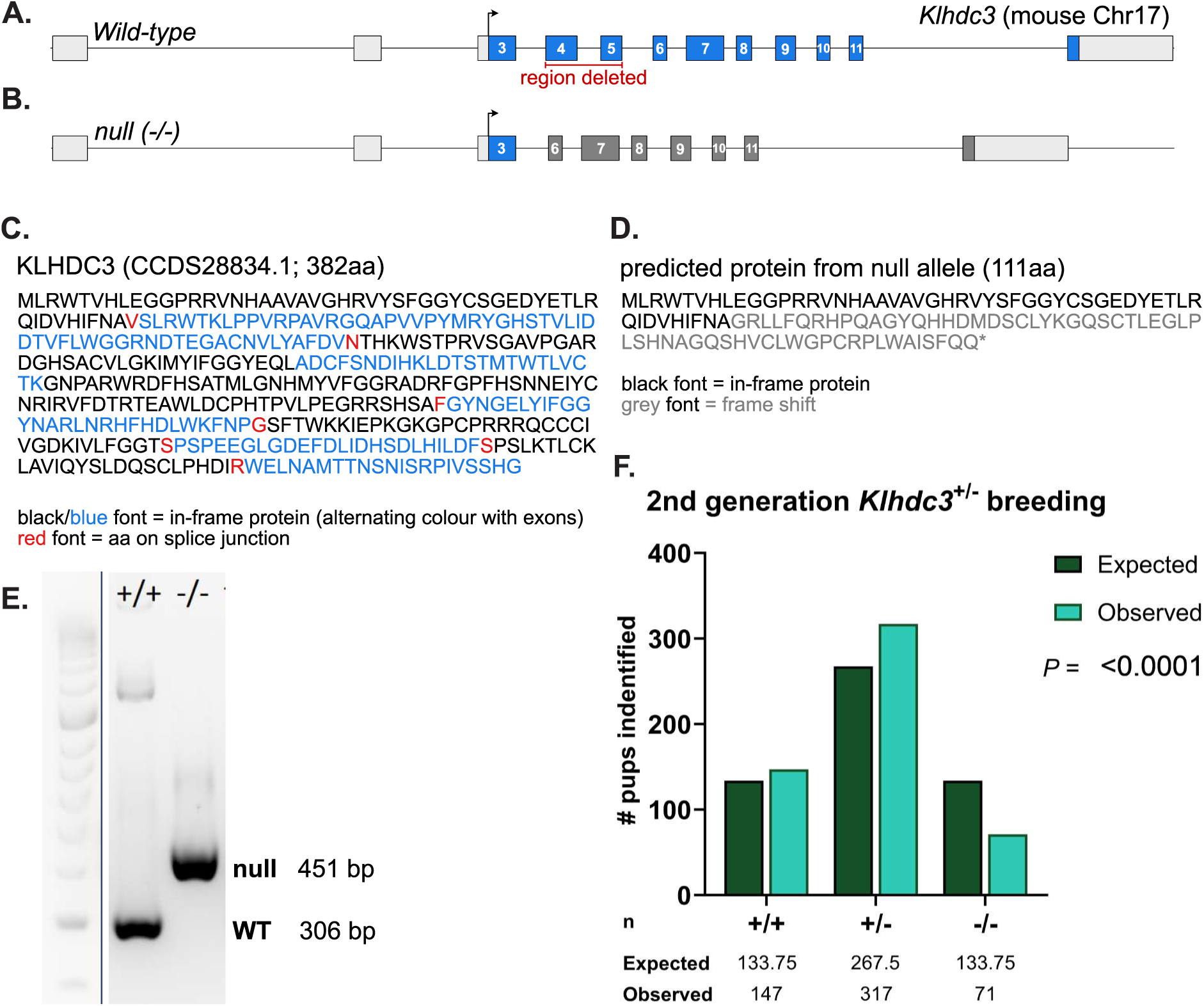
Generation of *Klhdc3* deficient mouse models. (A) Schematic of the wild type *Klhdc3* locus. (B) Germ-line null allele (*Klhdc3^-/-^*) identified during targeting to insert the LoxP elements. This line was bred separately from the conditional *Klhdc3^fl/fl^*allele. (C) The amino acid sequence of the WT KLHDC3 protein (D) The amino acid sequence of the predicted product of the deleted allele (expected to be the same for both the germ-line deficient and recombined LoxP flanked allele). (E) Representative genotyping from DNA of the indicated genotypes. (F) Genotype recovery of F2 generation pups born from the *Klhdc3^+/-^* x *Klhdc3^+/-^*crosses. Significant difference (p<0.0001) against the expected ratio was determined using Observed vs. Expected test in GraphPad Prism.

### *Klhdc3*-deficient mice are viable, runted and recovered at sub-Mendelian ratios

Due to the isolation of the germ-line *Klhdc3^+/-^* allele, we were able to establish breeding pairs of *Klhdc3^+/-^*mice and monitor the resultant litters. We examined the recovery of animals of all genotypes using DNA samples collected at day 7-10 after birth and by PCR identified multiple animals with homozygous *Klhdc3* deletion (Fig 1E). We were not able to confirm loss of KLHDC3 protein, as none of the commercial or custom-made antibodies we have tested were found to be specific. While the *Klhdc3*^-/-^ mice were viable, they were present at sub-Mendelian ratios with ∼50% of expected animals identified at day 7-10 sample collection (Fig 1F). Having recovered viable *Klhdc3*^-/-^ animals, we tested if they were fertile by attempting to breed male (n=2) and female (n=2) *Klhdc3*^-/-^ mice with *Klhdc3^+/-^*or WT mice. Albeit from a small sample size, we did not observed a confirmed pregnancy or viable offspring from the *Klhdc3*^-/-^ mice, indicating that the *Klhdc3*^-/-^ mice are most likely infertile. KLHDC3 is highly expressed in the testis in both human and mouse (Fagerberg *et al*, 2014; Yue *et al*, 2014), which may be relevant to the male infertility. It is unclear at present what accounts for the lack of viable pregnancies in the *Klhdc3*^-/-^ female mice.

We monitored the litters from the breeding pairs of *Klhdc3^+/-^*mice. Weights were recorded at weaning (day 19-29), with weaning age determined by animal facility staff according to management standard operating procedures and an assessment of suitability to be weaned independent of researchers and blind to genotype. At weaning, *Klhdc3^+/+^* and *Klhdc3^+/-^* mice had no significant difference in weight, while *Klhdc3^-/-^* mice had a significantly lower weight (∼70% of +/+ or +/-), regardless of the age at which they were weaned (Fig. 2A-B). The average weight of *Klhdc3*^-/-^ mice was 7.7g whereas wild type mice weighed an average of 10.2g and heterozygous mice 11.2g (Fig 2C).

**Figure 2.**
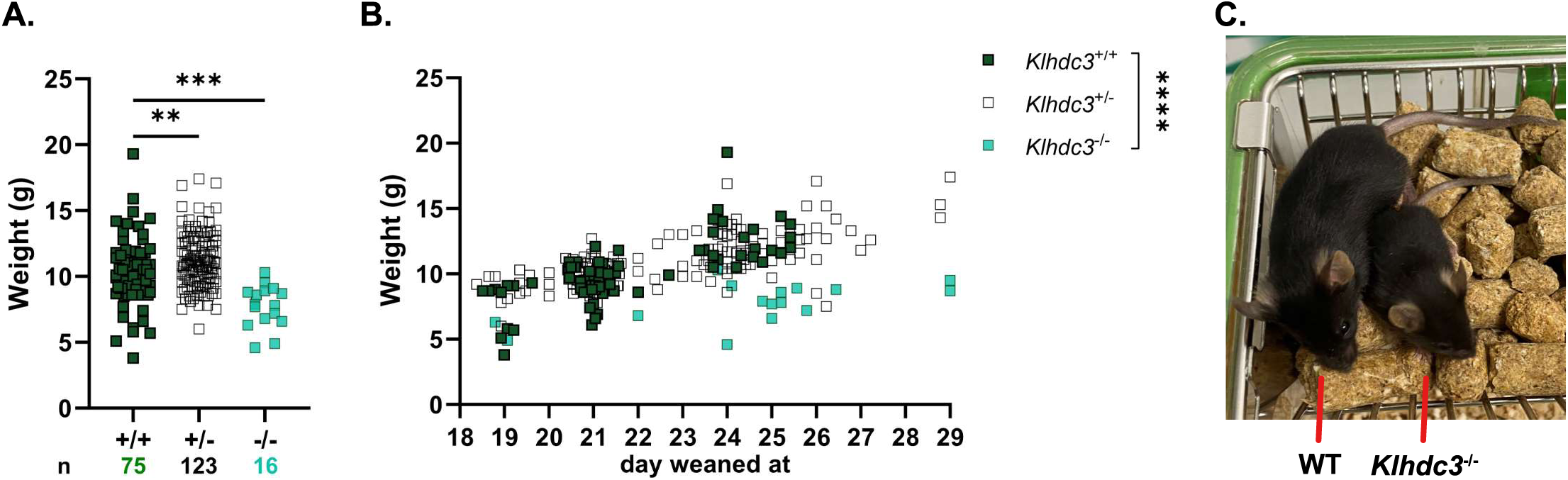
*Klhdc3* knockout mice have significantly lower weaning weights. (A) Mean weaning weights of indicated genotypes; Ordinary One-way ANOVA. (B) The weaning weights form panel (A) displayed according to the specific day of weaning (Two-way ANOVA; ns: p>0.05, *p≤ 0.05, **p≤ 0.01, ***p≤ 0.001, ****p≤ 0.0001). Weaning day was determined by animal facility staff based on SOP and assessment of suitability to be weaned independent of researchers and blind to genotype. *Klhdc3*^+/+^ n=75, *Klhdc3*^+/-^ n=123, *Klhdc3*^-/-^ n=16. (C) Photo of littermate pups show *Klhdc3*^+/+^ and *Klhdc3^-/-^*. Image at 23 days old.

### Adult *Klhdc3^-/-^* mice have reduced long-term survival

The *Klhdc3^-/-^* mice were smaller and had reduced weight, even as young adults (Fig 3A), although this became less pronounced as the *Klhdc3^-/-^* mice aged and appeared to gain weight and mass in a manner comparable to their littermates. The *Klhdc3^-/-^* mice that survived past the first month were often euthanised for ethical reasons due to illness or injury/wounds. Male *Klhdc3^-/-^* mice were frequently found to have a sore or wounded penis (Fig. 3B), whereas several female *Klhdc3^-/-^* mice had ulcer-like lesions on their abdomen. To understand the phenotype in more detail, one male and one female WT and *Klhdc3*^-/-^, respectively, from the same litter were sent for a histopathological analysis at 8 weeks of age (Fig 3A; Supplemental dataset 1). All animals were in good general condition, with both *Klhdc3*^-/-^ mice noted as smaller than the wild-type littermates (male *Klhdc3^-/-^* 17.7g vs. WT 24.4g and female *Klhdc3^-/-^*16.2g vs. WT 18.8g). The analysis assessed ∼25 tissues and organs (Supplemental dataset 1). The most notable findings from this analysis were that the male *Klhdc3^-/-^* had vacuolation/degeneration apparent in the heart and mild lymphocytic infiltrates in the dermis. The dermal infiltrates are of note, given that the aged *Klhdc3^-/-^* animals often developed skin lesions that ultimately required euthanasia. It is not at present clear if these are directly related observations. No pathological changes were observed in the testis. The female *Klhdc3^-/-^* mammary glands had a paucity of lactiferous ducts when compared to her littermate control. The uterus of the female *Klhdc3*^-/-^ was thin and delicate, and the ovaries were inconspicuous. The thinner endometrium may be relevant to the breeding failure of female *Klhdc3*^-/-^ mice. The full report of the histopathological findings can be found in Supplementary dataset 1.

**Figure 3.**
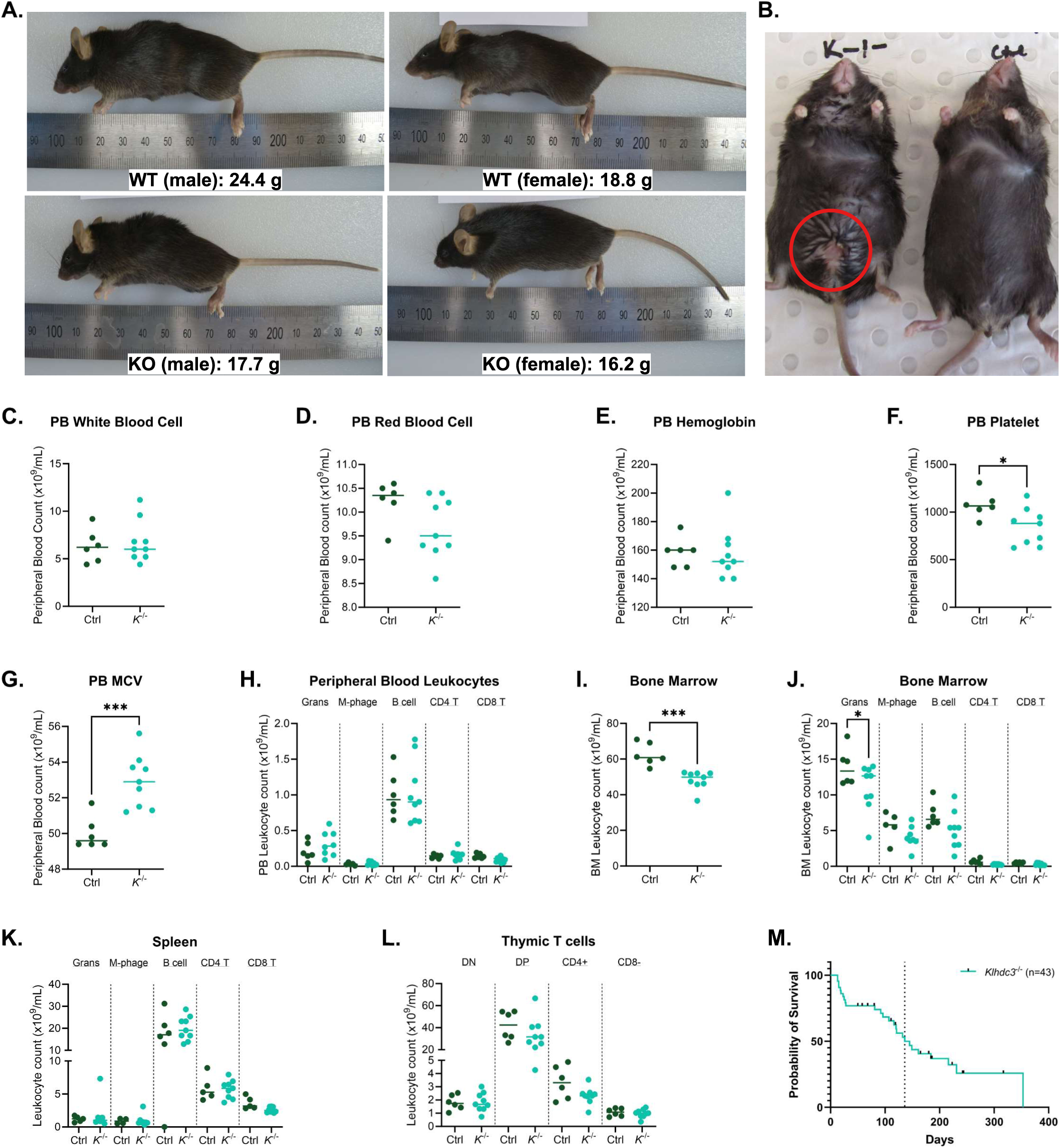
*Klhdc3^-/-^* mice have no apparent defects but show reduced survival rates. (A) Representative photos of littermate animals at 8 weeks old. (B) *Klhdc3^-/-^*mice develop skin lesions, which on male mice most often occur around the penis and on the lower abdomen. Male *Klhdc3^-/-^* at 242 days old, compared to co-housed control mouse. (C-G) Peripheral blood parameters: leukocyte count (C), red blood cell count (D), haemoglobin (E), platelet number (F) and red cell mean corpuscular volume (G) for *Klhdc3^+/+^* (control) and *Klhdc3^-/-^* animals. (H) Differential analysis of peripheral blood leukocytes: Grans: granulocytes, M-phage: macrophages, B220+ B cells and CD4+ or CD8+ T cells. (I) Bone marrow cellularity (per 2 femurs) and (J) lineage distribution per 2 femurs (as in H) for *Klhdc3^+/+^* (control) and *Klhdc3^-/-^* animals. (K) Spleen leukocyte lineage distribution (as in H). (L) Thymus T cell populations: DN: CD4 CD8 double negative, DP: CD4 CD8 double positive and CD4+ or CD8+ single positive cells. (M) Kaplan-Meier plot showing survival analysis of *Klhdc3*^-/-^ mice. n=43, median survival=136 (as indicated by dotted line).

We further assessed the haematological parameters of the *Klhdc3^-/-^*mice as they were identified as requiring euthanasia primarily for skin lesions/wounds. Co-housed littermates were assessed as references in each case. In the peripheral blood, there were only subtle overall differences compared to littermate controls (Fig 3C-H). While total leukocyte numbers were normal (Fig 3C), there was a reduced platelet count (Fig 3F) and increased mean corpuscular volume of the red blood cells (Fig 3G). There was no difference in the immune cell populations of the myeloid or lymphoid lineage in the peripheral blood in the *Klhdc3*^-/-^ mice (Fig 3H). There was a reduction in the overall cell number in the femurs of the *Klhdc3*^-/-^ mice (Fig 3I), however this could reflect the reduced size of the *Klhdc3^-/-^* mice compared to their controls, and a slight but significant reduction in the bone marrow granulocytes (Fig 3J). There was no difference in the number of cells of each lineage in the spleen (Fig 3K) or thymus (Fig 3L).

We then monitored the *Klhdc3^-/-^* mice as they aged into adult animals and observed reduced viability of the *Klhdc3^-/-^*mice (Fig 3M). Approximately 25% of the *Klhdc3^-/-^*pups identified at day 7-10 were found dead within the first month after birth without specific cause of death. The surviving mice appeared normal and healthy until the age of 3 months. From then on mice were regularly found to be moribund or to have lesions/wounds that required euthanasia. Post-mortem examination however, failed to reveal any evidence of causative disease. Only a few individuals survived to ∼1 year of age resulting in an overall median survival of 136 days (Fig 3M).

### *Klhdc3^-/-^* mice become obese with age

As we observed the mice with aging, it became apparent that the *Klhdc3^-/-^* mice were becoming obese (Fig 4A). This occurred in settings of mixed housing and *ad libitum* chow, where littermates/age matched animals were co-housed but the control animals did not become obese. Autopsy analysis indicated that the older *Klhdc3^-/-^* mice had excessive adipose tissue, macroscopically visible as subcutaneous deposits and in both the peritoneal and chest cavity (Fig 4A). Based on this we re-assessed the samples that were used for histopathological assessment and could observe an increased adipocyte layer in the skin and mammary tissue of the *Klhdc3^-/-^*animals compared to their littermate sex matched controls (Fig 4B). This was apparent in both male and female 8-week-old *Klhdc3^-/-^* mice compared to WT littermates, despite these 8-week-old mice being of lower body weight. This occurred in settings of mixed housing and *ad libitum* chow, where littermates/age-matched animals were co-housed, but the control animals did not become obese.

**Figure 4.**
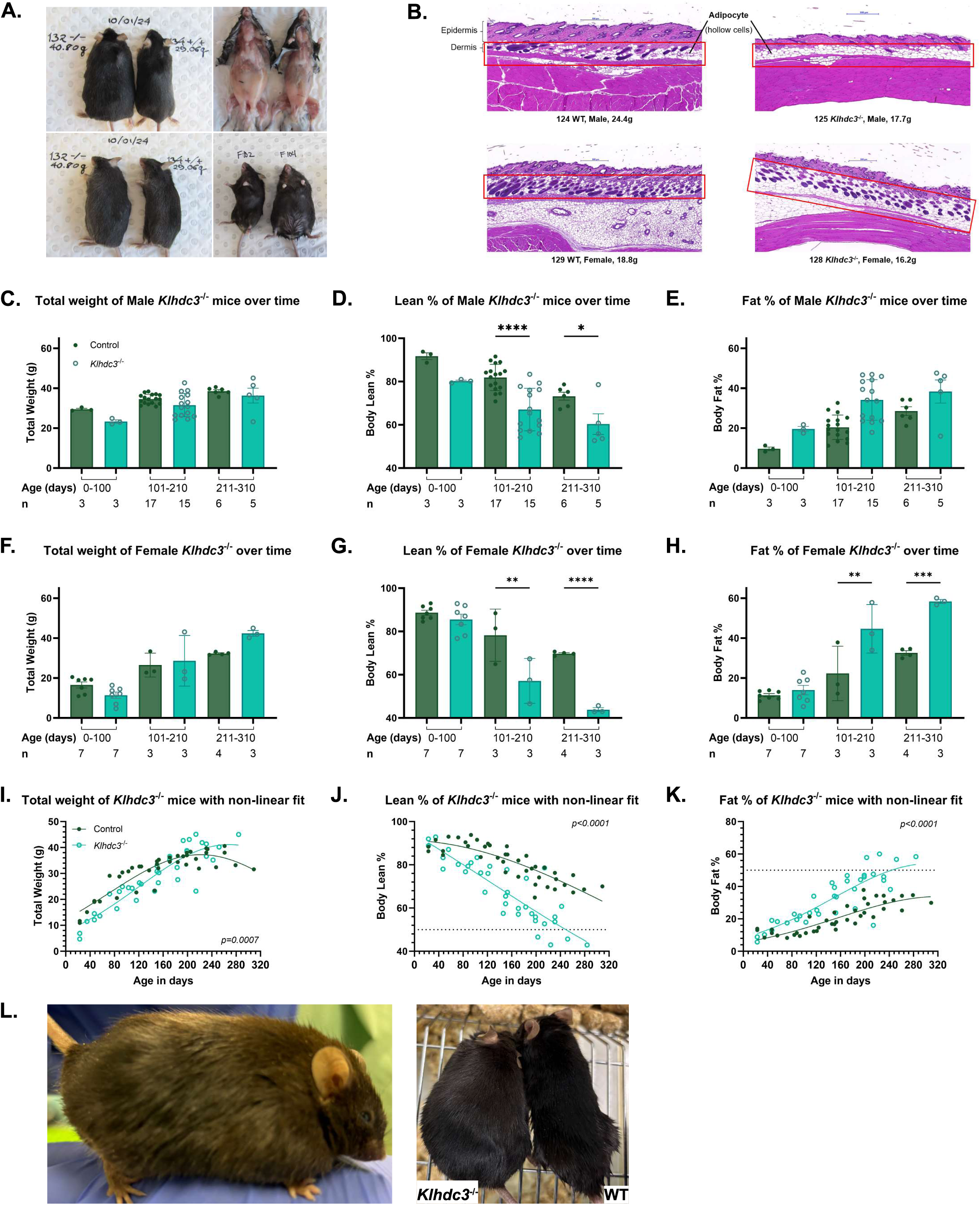
*Klhdc3^-/-^* mice develop age-related obesity. (A) *Klhdc3^-/-^*mice have excessive subcutaneous and visceral fat compared to co-housed littermate controls. Male Klhdc3 KO (#132) and male WT (#134) at 184 days of age. Female Klhdc3 WT (#102) and female Klhdc3 KO (#104) at 199 days of age. (B) Representative histological analysis of the skin of the indicated genotypes and sexes showing increased adipocyte numbers in *Klhdc3^-/-^* mice. (C) Total body weight of male mice grouped by age and genotype as indicated. Animals were longitudinally assessed. (D) Percentage lean mass and (E) Percentage body fat of male mice analysed using EchoMRI grouped by age and genotype as indicated. Animals were longitudinally assessed. (F) Total body weight of female mice grouped by age and genotype as indicated. Animals were longitudinally assessed. (G) Percentage lean mass and (H) Percentage body fat of female mice analysed using EchoMRI grouped by age and genotype as indicated. Animals were longitudinally assessed. (I) Total body weight (p=0.0007), (J) Percentage lean mass (p<0.0001) and (K) Percentage fat mass (p<0.0001) of *Klhdc3*^+/+^ and *Klhdc3*^-/-^ mice. Genotypes as indicated and both sexes pooled. Animals were longitudinally assessed. The line of best fit is indicated for each genotype. The dashed line in panels J and K represent the 50^th^ percentile. (L) Female *Klhdc3*^-/-^ mouse at 224 days old, measured to have 60.05% body fat content, in comparison to co-housed WT control at 219 days old and 34.29% body fat content. Bar graphs were tested using Ordinary one-way ANOVA, with sample size indicated below the graph. Non-linear curve fits in panels I-K have been compared using F test (ns: p >0.05, * p ≤0.05, ** p ≤0.01, *** p ≤0.001, **** p ≤0.0001).

We then assessed a cohort longitudinally using EchoMRI to allow non-invasive measurement of fat to lean body mass ratio. We measured body weight, lean mass and fat mass parameters over time, with the data analysed by age groups for males and females separately (Fig 4C-4H), and by specific age for both sexes combined (Fig 4I-4K). At around 40-45 days old, both male and female *Klhdc3^-/-^* mice were runty and had a lower total body weight compared to control littermates (Fig 4C, 4F, FI). As the mice aged the females developed a significantly higher proportion of body fat as a percentage of total body mass compared to controls (Fig 4E, 4H, 4K). This increase in fat accrued at the expense of lean mass, which by body weight percentage was found to be compromised from 45 days of age onward (Fig. 4D, 4G, 4J). The difference between WT and *Klhdc3^-/-^* mice (sex pooled data) in the proportion of fat to lean body mass percentage diverged around the 80-day age mark and became progressively more pronounced with additional age (Fig 4I-4K). At ∼200 days of age, the total body weight of *Klhdc3^-/-^* mice began to approximate the weight of the control mice (Fig. 4K). Strikingly, at this timepoint and from here on till the last assessment of the *Klhdc3^-/-^*mice at 290 days, ∼50% of the total mass is derived from fat mass (Fig 4K-4L). These observations indicate that KLHDC3 plays an important role *in vivo* in restricting adiposity and fat mass *in vivo*.

### *In vitro* analysis of KLHDC3-deficient cells

Mouse embryonic fibroblasts (MEFs) were prepared from embryos collected at embryonic day 12.5 and E14.5. The MEFs were assessed both as primary and immortalised cell lines through the shRNA-mediated knock-down of *p53* (Dickins *et al*, 2005; Mutsaers *et al*, 2013; Ng *et al*, 2015). The primary cell lines were passaged alongside the counterpart immortalised cell lines to obtain growth curves for both control and *Klhdc3^-/-^* fibroblasts (Fig 5A-5B). There was no significant difference in either proliferation rates or viability between the *Klhdc3^-/-^* cells and *Klhdc3* WT or heterozygous controls, either as primary or immortalised lines (Fig 5A-5B). To assess the fraction of actively replicating cells the immortalised MEFs were pulse labelled with EdU. No significant difference was observed in the proportion of *Klhdc3*^-/-^ MEFs undergoing DNA replication (S-phase) compared to the wild type or heterozygous MEFs (Fig. 5C). These results collectively indicate that the deletion of *Klhdc3* is tolerated *in vitro* and does not alter cell proliferation.

**Figure 5.**
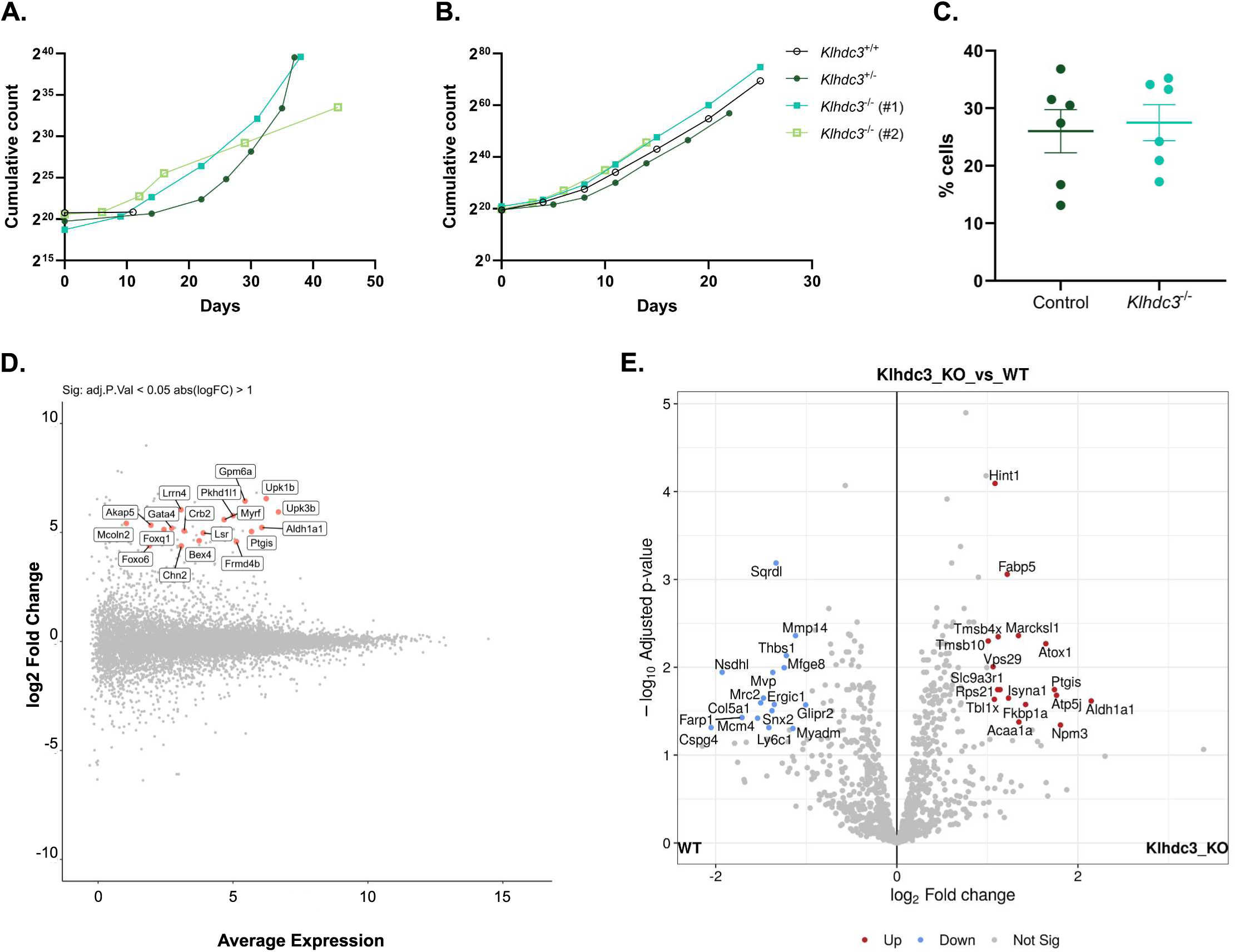
Transcriptomic and proteomic analysis of *Klhdc3*^-/-^ deficient mouse embryonic fibroblasts. (A) Proliferation curves of primary E14.5 mouse embryonic fibroblasts (MEFs) of the indicated genotype. (B) Proliferation curves of immortalised E14.5 mouse embryonic fibroblasts (MEFs) of the indicated genotype. (C) Asynchronous immortalised MEFS of the indicated genotypes were assessed for cell cycle distribution; S-phase proportion assessed using EdU incorporation. An unpaired t-test shows no significance between the control and *Klhdc3*^-/-^ cell lines. Each control data is an independent cell line. *Klhdc3^-/-^*has 5 independent cell lines, one of which had been replicated. (D) Transcriptomic analysis of immortalised control and *Klhdc3*^-/-^ MEFs with differentially expressed genes indicated in red and labelled; n=3 per genotype. (E) Proteomic analysis of differentially expressed proteins in the immortalised control and *Klhdc3*^-/-^ MEFs; data displayed as a volcano plot of the adjusted p-value against the fold change; n=6 per genotype. Both transcriptomic and proteomic results have been filtered to those with an adjusted p <0.05 and abs(log_2_FC) >1.

We then assessed the transcriptome of both *Klhdc3*^-/-^ MEFs (Fig 5D; Supplemental dataset 2) and *Klhdc3-*deficient immortalised myeloid cells (generated through CRISPR/Cas9 targeting; Supplemental dataset 3) to understand the transcriptional consequences of loss of KLHDC3. Only one transcript (*Padi4*) was differentially expressed between *Klhdc3^-/-^* and *Klhdc3^+/+^*myeloid cells, so they were not further analysed (Buco *et al*., 2025). In the MEFs, differentially expressed genes (DEGs) were filtered to include only those with an absolute Log_2_ fold change of ≥1 and a false discovery rate of ≤0.05 (Supplementary dataset 2). Consistent with a primary function of KLHDC3 in post-translational regulation of proteins, there were only a very small number of changes apparent in the transcriptome and all of these showed increased expression in the *Klhdc3*^-/-^ MEFs. Of the 18 upregulated transcripts, pathway enrichment analysis identified two involved in lipid metabolic pathways, namely *Aldh1a1* and *Ptgis*. These limited changes overall, indicate that loss of KLHDC3 does not substantially alter the transcriptome.

Given the proposed role of KLHDC3 in post-translational regulation via the DesCEND pathway, we conducted a proteomic analysis to identify differentially expressed proteins in the *Klhdc3*^-/-^ MEFs (Fig 5E). For this analysis, we utilised the same MEFs as were used for the RNA sequencing experiment (Fig 5D) to allow a comparison between the transcriptome and proteome profiling. Protein samples were collected and were digested into peptides and separated using liquid chromatography before conducting mass spectrometry. A total of 1146 proteins were identified. The differentially expressed proteins (adj. p-value <0.05, FC >1) are shown in a volcano plot (Fig. 5E, Supplemental dataset 4). Pathway analysis of the proteins that were increased or decreased in abundance in the *Klhdc3*^-/-^ MEFs identified that 5 of the 17 upregulated and 1 of the 16 downregulated proteins were involved in the lipid metabolic process including ALDH1A1, PTGIS, ACAA1A, ISYNA1, FABP5 and NSDHL, with *Aldh1a1* and *Ptgis* also showing increased transcript expression (Caldas & Herman, 2003; Carbonetti *et al*, 2019; Sasaki *et al*, 2021; Wang *et al*, 2020; Wang *et al*, 2021; Xu *et al*, 2022; Yang *et al*, 2017; Zhang *et al*, 2020; Ziouzenkova *et al*, 2007). This was of particular interest considering the obese phenotype we observed in the *Klhdc3*^-/-^ mice. The literatures correlating the genes of interest to a function in lipid metabolism support their role in obesity, apart from ACAA1 whose knockdown was shown to increase lipid accumulation in preadipocyte differentiation from sheep (Wang *et al*., 2021), whereas we see an increase in expression in our *Klhdc3^-/-^* samples. Due to KLHDC3 having sequence-specific substrate recognition, the results were further filtered to those predicted to contain a terminal glycine residue. From this analysis, only HINT1 fulfilled the criteria, which coincidentally was the most significantly differentially expressed protein based on p-value from the proteomics (Fig 5E). Murine HINT1 has a C-terminal RQMNWPPG* sequence and was increased >2-fold in expression in the *Klhdc3*^-/-^ MEFs. Therefore, HINT1 was further investigated as a potential target of KLHDC3.

### Validation of HINT1 as a target of KLHDC3

To determine if HINT1 was a target of KLHDC3, we established HINT1 global protein stability (GPS) reporter assays (Fig 6A-6D). In this assay a reporter is generated where two fluorescent proteins are expressed: a DsRed that is expressed from the CMV promoter, followed by an IRES, and an eGFP fused to the protein of interest (Koren *et al*., 2018; Lin *et al*., 2018; Yen & Elledge, 2008; Yen *et al*, 2008). The level of DsRed expression is stable while the level of eGFP depends on the stability of the fused protein. Thus, the ratio of DsRed to GFP can be used as a reflection of stability of the protein of interest, in this case HINT1. GPS reporter constructs were designed using HINT1 from both mouse and human. Human and mouse HINT1 are 94% conserved overall and the C-terminal motif is RQMxWPPG*, where x is N in mouse and H in human.

**Figure 6.**
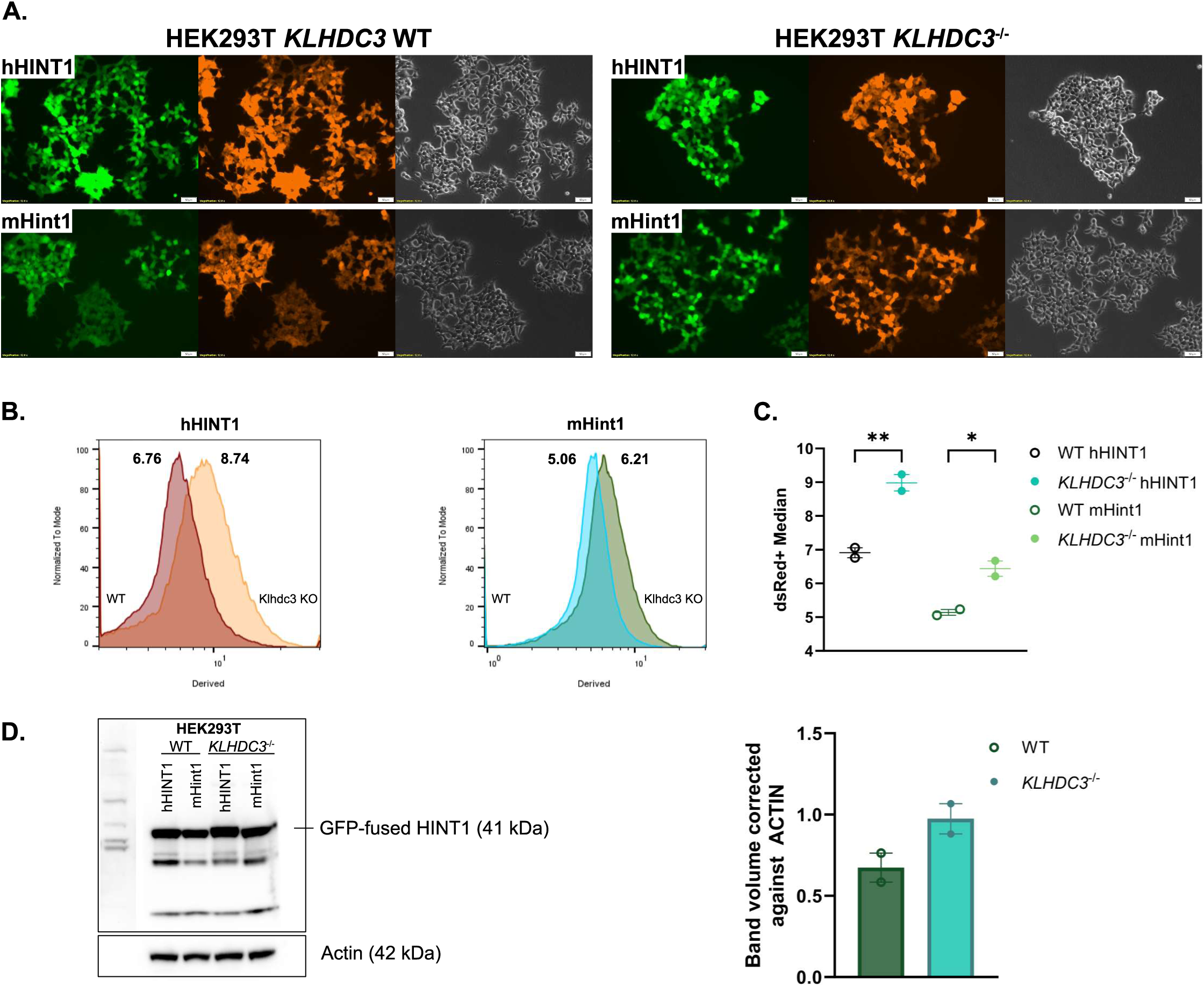
Validation of HINT1 as a new target of KLHDC3. (A) Human *HINT1* (hHINT1) or murine *Hint1* (mHint1) cDNAs were cloned into the GPS-reporter plasmid. In this plasmid DsRed is constitutively expressed but GFP expression is determined by the stability of the fused HINT1 protein. This reporter was expressed in KLHDC3 WT HEK293T cells or *KLHDC3^-/-^* HEK293T cells as indicated. Fluorescence signal was detected by live cell fluorescent imaging. Scale bar represents 50μm. (B) Flow cytometric quantitation of the GFP expression levels of human or murine HINT1 fusion proteins in KLHDC3 WT or KO HEK293T cells. Data shown as the GFP/DsRed derived median with a higher value indicating increased stability. (C) Quantitation of the GFP/DsRed derived median from human and mouse HINT1 in the WT and *KLHDC3^-/-^* HEK293T cells shown in B. Data from replicate experiments. (D) Western blot from extracted GFP/DsRed human and mouse HINT1 proteins. Quantification plots the volume of the band (intensity) corrected against ACTIN. Not significant using t-test.

We generated HEK293T cells that were either WT or deficient in KLHDC3 (*KLHDC3^-/-^*) (Buco *et al*., 2025). The GPS reporter constructs were transfected into the HEK293T cell lines and the DsRed and GFP signals assessed by both fluorescent microscopy and flow cytometry (Fig 6A-6C). These analyses demonstrated an increased GFP signal in *KLHDC3^-/-^* HEK293T compared to *KLHDC3* WT HEK293T cell lines, reflecting the KLHDC3 dependent stability of HINT1 (Fig 6C). This was the case for both human and mouse HINT1.

In addition to the GPS reporter assay, western blotting was undertaken of proteins extracted from the GFP-HINT1 transduced cell lines using a HINT1 antibody (Fig 6D). Consistent with the GPS reporter assay, WT HINT1 from both human and mouse was more highly expressed in *KLHDC3^-/-^* 293T than in WT HEK293T cell lines (Fig. 6D). These results support the conclusion that HINT1 is a physiological target of KLHDC3.

## Discussion

Here we report the development and characterisation of the first *Klhdc3*-deficient mouse model. This model was established to understand the physiological functions and to assist in the identification of substrates of KLHDC3, a CUL2 E3 ligase receptor of the DesCEND pathway. We generated *Klhdc3^-/-^*animals and demonstrated that these are born smaller and at sub-Mendelian numbers (∼50% of expected). The cause of the pre-natal developmental lethality is not clear from the available data, but is consistent with the analysis of human variants, where KLHDC3 is intolerant to loss of function variants (gnomAD browser) (Chen *et al*, 2024; Karczewski *et al*, 2020). The KLHDC3-deficient animals had increased post-natal lethality. A broad-based histopathological assessment of 8-week-old male and female *Klhdc3^-/-^* animals did not identify any specific abnormalities. The viable *Klhdc3^-/-^* animals are smaller at weaning but become obese in adulthood. To our surprise we found a high body fat percentage already present at an early age, even when the *Klhdc3^-/-^* animals are runted relative to littermate controls. It must be noted however, that we did not observe any of these phenotypes in heterozygous littermate controls. To date mutations in KLHDC3 have not been reported in association with obesity in humans (Loos & Yeo, 2022). Querying the Musculoskeletal Knowledge Portal (https://mskkp.org; search term: KLHDC3) for human genetic evidence for associations of KLHDC3 variants with human phenotypes there are moderate associations a range of traits overlapping with the murine *Klhdc3^-/-^* phenotype, including with waist-hip ratio, height, a range of lipid traits and appendicular lean body mass (Dornbos *et al*, 2022).

We tested breeding of a small number of the *Klhdc3^-/-^*animals, both males and females, and never observed either a confirmed pregnancy (by palpation) or any pups born from these crosses (all to *Klhdc3^+/-^*or C57BL6/J animals), indicating that both sexes are most likely infertile. When originally identified, *Klhdc3* was shown to be expressed from a testis-specific bidirectional promoter with male enhanced antigen (MEA1) and a third protein PP2R5D that form an overlapping gene complex (PMP- complex) (Ohinata *et al*, 2003b). This is conserved across human and mouse. In mouse testis *Klhdc3* is expressed in a stage-specific manner with expression beginning in mid-spermatogenesis and the protein predominantly localizing to the cytoplasm and meiotic chromatin of pachytene spermatocytes, suggesting an involvement in the process of meiotic recombination (Ohinata *et al*., 2003a). Single cell analysis of human samples indicated late spermatid specific high expression of KLHDC3 (https://proteinatlas.org, (Uhlen *et al*, 2015) and some studies have suggested a correlation between downregulation of *KLHDC3* and globozoospermia or azoospermia (Cheung *et al*, 2021). The mouse ENCODE transcriptome data demonstrates that while *Klhdc3* is most highly expressed in the testis (mean RPKM ∼250), it is broadly expressed at significant levels (RPKM>25) in the majority of tissues assessed (Yue *et al*., 2014).

Only limited information is available on the *in vivo* functions of other C-degron pathway and KLHDC members. Homozygous *Klhdc2*-deficient mice are reported as non-viable (Dickinson *et al*, 2016). Interestingly, given the phenotype we observed with our *Klhdc3^-/-^* animals, female *Klhdc2^+/-^* animals were reported to have increased adiposity (∼135% of the control cohort) as part of a high-throughput phenotyping program (*Klhdc2^tm1b(EUCOMM)Hmgu^* allele; MRC Harwell data from IMPC portal; p=1.94x10^-6^). Another E3 ligase of the C-degron pathway, CUL2-APPBP2, regulates the stability of PRDM16 and thereby beige fat biogenesis (Wang *et al*, 2022). The result is, however, the opposite of what we report here in the *Klhdc3^-/-^* where we observed increased adiposity. More puzzling given our phenotype and the *Klhdc2^+/-^*finding, an adipocyte-specific CUL2 deletion, which engages APPB2 as well as KLHDC2 and KLHDC3, protected against diet-induced obesity (Wang *et al*., 2022). *Klhdc10^-/-^*mice are viable (Yamaguchi *et al*, 2016). The *Klhdc10*-deficient mice were reported to have an altered response to inflammatory stimuli and the results suggested a function in necroptosis (Yamaguchi *et al*., 2016). This analysis, albeit of only a limited number of the KLHDC members to date, emphasises that although very similar in structure and substrate binding mechanism, these substrate receptors have different targets and function in different pathways *in vivo*. Assessing the IMPC database for phenotypes associated with increased adipose tissue highlights roles of known genes (e.g. *Lepr*, *Ghrh*, *Gh*) and many not previously associated with increased adiposity among the 4.45% of gene mutations that have this phenotype (366 genes with a significant difference of 8222 tested genes; mammalian phenotype browser term MP:0010024). Our analysis of the *Klhdc3^-/-^* together with previously reported data suggest a broader function for the C-degron pathway in adipose tissue homeostasis.

At a cellular level we did not observe any significant differences in cell viability, proliferation or cell cycle of mouse embryonic fibroblasts. We also assessed immortalised myeloid cell lines that we engineered to be KLHDC3-deficient with CRISPR/Cas9, and these were also not discernibly different to WT cells in terms of proliferation kinetics, viability or transcriptome (Buco *et al*., 2025). Similarly, we observed largely normal haematopoiesis in the adult animals even when requiring euthanasia due to skin lesions or degree of obesity. Collectively, these results together with the *in vivo* phenotype we observed, would indicate that the function of KLHDC3 is cell type and developmental stage dependent. Our analysis using RNA-seq demonstrates that loss of KLHDC3 only resulted in subtle effects on the transcriptome, consistent with the post-translational function of KLHDC3. At the protein level we could identify a range of proteins that were differentially expressed following the loss of KLHDC3, with a significant proportion of these linked to lipid metabolism. Only one of the upregulated proteins, however, is likely to be a direct substrate of KLHDC3.

The protein that had the most significantly altered expression in the *Klhdc3*^-/-^ MEFs, as well as a protein containing a C-terminal glycine, was HINT1. Other proteins previously identified and validated *in vitro* were either not amongst the 1146 isolated proteins (PPP1R15A, USP49, TSPYL and TCAP) or contain a different C-degron in mouse (p19^ARF^) (Koren *et al*., 2018; Lin *et al*., 2018; Yeh *et al*., 2021). HINT1 (histidine triad nucleotide binding protein 1, also known as HINT; NMAN; PKCI-1; PRKCNH1) is an adenosine 5’-monophosphoramide hydrolase (Brenner, 2002; Zhou *et al*, 2013). Loss-of-function variants in HINT1 have been reported as a cause autosomal recessive neuromyotonia and axonal neuropathy (Kontogeorgiou *et al*, 2021; Zhao *et al*, 2014; Zimon *et al*, 2012). Here we find that loss of KLHDC3 leads to increased levels of HINT1. The consequences of increased HINT1 on the phenotypes we observed in the *Klhdc3*^-/-^ animals *in vivo* are not clear. There are increasing reports of the consequences of increased HINT1 in a range a contexts, including cancer, neuronal cells and cardiac function (Kang *et al*, 2025; Wang *et al*, 2023; Zhang *et al*, 2025). We expect that HINT1 is only one of the KLHDC3 targets that become deregulated and that a more comprehensive assessment of tissues, such as adipocytes or their precursor cells, would be informative to identify potential KLHDC3 substrates that directly mediate the adipose tissue defects we observed. An alternative interpretation is that the adipose tissue is expanding due to a failure in the lean tissue mass to achieve its appropriate set point. This seems less likely, however, as we could observe increased fat mass even in runty animals early in life and this then expands to be beyond 50% of the total animal mass by ∼220 days of age.

In summary, we describe here the development and characterisation of a *Klhdc3*-deficient mouse model. This model demonstrates a previously unappreciated role for the DesCEND pathway, and KLHDC3 specifically, in regulating normal development and homeostasis. We demonstrate a key homeostatic function for KLHDC3 in restricting post-natal adiposity and body fat accumulation, with adult *Klhdc3^-/-^* mice having more than 50% of their body mass derived from fat. We further identified that HINT1 is a new physiological native protein target for KLHDC3. The findings presented here extend our understanding of the DesCEND pathway to *in vivo* models and identify KLHDC3 as a critical regulator of organismal homeostasis and longevity.

## Materials and Methods

### Ethics Statement

All animal experiments were performed in compliance with ‘The Prevention of Cruelty to Animals Act’ (1986), the associated Regulations and the National Health and Medical Research Council ‘Australian Code for the Care and Use of Animals for Scientific Purposes (2013). The procedures were approved by the Animal Ethics Committee (AEC) at St. Vincent’s Hospital, Melbourne (AEC Reference #019/21-r3).

### Generation of Klhdc3 mouse models

*Klhdc3* conditional (*Klhdc3^fl/fl^*) mice were generated by the Monash Genome Modification platform by CRISPR/Cas9 mediated insertion of loxP elements flanking *Klhdc3* exons 4 and 5 (Buco *et al*., 2025). A ssDNA repair template was prepared for the required sequence to insert. Cas9 protein (80ng/μl; Integrated DNA Technologies (IDT)) was incubated with sgRNA (80ng/μl; crRNA and tracrRNA; IDTDna) to generate RNP. sgRNA used for targeting: sgRNA1: 5’ TGGTGGTCTCTGCGGCAAGG 3’; sgRNA2: 5’ TGCCACTTAAATGGTGTCAA 3’.The ssDNA repair template containing the LoxP flanked region was generated using the Guide-it Long ssDNA Production System (Takara) according to the manufacturer’s instructions. The RNP and the ssDNA repair template (50ng/ml) were microinjected into C57BL/6J zygotes at the pronuclei stage. Microinjected zygotes were transferred into the uterus of pseudo pregnant F1 females. Correct insertion was screened by genomic DNA PCR and then digital droplet PCR (Bio-Rad). Correctly targeted males were identified and bred to C57BL/6J females to ensure germ-line transmission. The *Klhdc3^fl/fl^*allele has not been used for the experiments described in this study (Buco *et al*., 2025). During screening of progeny, a male with a deletion encompassing the loxP flanked region was identified (referred to from here on as *Klhdc3^+/-^*). This allele resulted from a fusion between the sgRNA induced breaks generating a 676bp deletion that removed exon 4 and 5 of *Klhdc3*. The male founder was bred to C57BL/6J females to ensure germ-line transmission.

All lines were confirmed to have the expected inserted elements/mutations by Sanger sequencing of the loxP elements and the deletion event, respectively. Second generation animals were then inbred to establish mice of the indicated genotypes. All animals were housed at the Bioresources Centre (BRC) located at St. Vincent’s Hospital, Melbourne. Mice were maintained and bred in microisolators under specific pathogen-free conditions with autoclaved food and acidified water provided *ad libitum*.

### Genotyping

DNA samples were extracted from either cultured cells or mouse tissues. The primers used for the detection of the alleles in Klhdc3, together with the expected product sizes, are as follows: WT: 5’-GCATGGTGGAACAAGAGACTTT-3’; Floxed: 5’-GCCAAGAGCAGAGAGATGGG-3’; Deleted allele (germ-line or recombined): 5’-CTGTGGCCCAGGGGATGTTA-3’; Product size: WT = 306bp; floxed = 386bp; deleted=451 (germ-line) or 552bp (recombination of the floxed allele).

### Body Composition Analysis

Live mice were weighed and placed in cylindrical tubes body composition assessment by EchoMRI (EchoMRI LLC) as described by the manufacturer.

### Blood and Tissue Analysis

Peripheral blood (PB) samples of mice were obtained through retro-orbital blood collection into BD microtainer tubes. Mice were euthanised through cervical dislocation or CO_2_ asphyxiation. The spleen, thymus, and femurs were collected, and single cell suspensions prepared in FACS buffer (PBS, 1% FBS). The cell suspensions from BM, spleen and thymus were filtered using a 40µm cell strainer. For whole blood cell counts, PB was analysed on a Sysmex KX21 (Roche Diagnostics) haematological analyser. For flow cytometric analysis, erythrocytes were removed from the PB using RBC lysis buffer (150mM NH_4_Cl, 10mM KHCO_3_, 0.1mM Na_2_EDTA, pH 7.3). Up to 1x10^6^ cells, were stained with fluorophore-conjugated antibodies and analysed on the BD LSRII Fortessa machine (BD Bioscience). Compensation and data acquisition were performed using the CellDiva software (BD Bioscience) and analysis using FlowJo software Version 9 or 10.0 (Treestar). All flow antibodies were purchased from ThermoFisher Scientific: anti-murine Gr1-FITC (Cat# 11-5931-85), CD11b/Mac1-APC-eF780 (Cat# 47-0112-82), B220-APC (Cat# 17-0452-83), CD4-eF450 (Cat# 48-0042-82), CD8a-PerCP-Cy5.5 (Cat# 45-0081-82), and TCRβ-PE (Cat# 12-5961-83). The samples were analysed on the BDLSRII Fortessa machine (BD Bioscience) and analysed using FlowJo software Version 9 or 10.0 (Treestar).

### Mouse embryonic fibroblasts cell lines

Females were timed-mated with males and E12.5 or E14.5 embryos were collected. DNA was isolated from a portion of the head for genotyping. The body was minced in 1mL of 0.025% trypsin and incubated at 37°C for 30-45 minutes. The tissue was then gently dissociated with a 1 mL pipette tip and diluted with a total of 10mL of media (DMEM, 10% FBS (not heat-inactivated; Bovogen Aust), 1% Glutamax and 1% Penicillin-Streptomycin; medias and additives from Gibco). Cells were left to grow to confluency under hypoxic conditions (5% CO_2_/5% O_2_/balance N_2_; Billups-Rothenberg modular hypoxic chamber) at 37°C. Once established the cells were cultured in standard 5% CO_2_ conditions at 37°C. Immortalisation of the cells was performed using retrovirally delivered shRNA against p53 (LMP-sh*Tp53*.1224) as previously described (Liang *et al*, 2023; Mutsaers *et al*., 2013).

### Cell culture and proliferation analysis

Cells were maintained in MEF medium in a controlled environment of 5% CO_2_ and at 37°C. Adherent cells were passaged by trypsinization upon reaching ∼70% confluence (every 3-14 days for primary cell lines, 2-5 days for immortalised cell lines). Cells were counted on a Countess™ II FL Automated Cell Counter (Thermo) using trypan blue as viability stain and re-seeded at the desired cell density in pre-warmed media.

### Cell cycle analysis

Cells were labelled by the addition of 10µM EdU to the medium followed by incubation at 37°C for 1hr. Cells were then collected by trypsinization, washed and resuspended in 100µL DPBS. Cells were fixed by adding 100µL 8% PFA and incubated for 15 min at room temperature. Cells were then washed twice with PBS/BSA (1%) and resuspended in 100-200µL of PBS/BSA (1%). Fixed cells were stained using AlexaFluor647-Azide and the Click-iT EdU Flow Cytometry Assay Kit (Thermo Scientific). Cells were resuspended in 100µL of DAPI (1µg/mL) and analysed on the BDLSRII Fortessa machine (BD Bioscience). Compensation and data acquisition were performed using the Cell Diva software and analysis using FlowJo software.

### RNA-sequencing

*Klhdc3*^+/+^ and *Klhdc3*^-/-^ MEFs were suspended in RLT buffer (Qiagen) and stored at-80°C until processed. RNA was purified using the RNeasy Mini Kit (Qiagen) as directed by the manufacturer. RNA quality was assessed using the RNA ScreenTape Assay for TapeStation Systems (Agilent). RNA (500ng per sample) was used to generate sequencing compatible libraries using the BRB-seq protocol (Alithea Genomics) (Alpern *et al*, 2019) as outlined by the manufacturer. The library was sequenced as PE150 by Novogene (Singapore) on the Illumina platform.

Raw reads were trimmed with fastp (Chen, 2023). BRB-Seq libraries were then demultiplexed using BRB-seqTools v1.6.1 (Alpern *et al*., 2019) and aligned using STAR v2.7.11a (Dobin *et al*, 2013) in quantMode to get GeneCounts (genome: mm10, gencode annotation: m22)

Differential gene expression analysis was then performed using the Degust analysis tool (Powell, 2019). Briefly, samples were normalized using the TMM method, the data was transformed to logCPM (moderated log counts per million). Genes were only considered with count >10 and CPM > 1 in at least three samples of a given genotype. Differential expression analysis was performed using the edgeR Quasi-Likelihood method (Chen *et al*, 2025).

### Mass spectrometry

Mouse embryonic fibroblasts were obtained from *Klhdc3*^+/+^ and *Klhdc3*^-/-^ embryos, grown to ∼70-80% confluency and washed and harvested. The proteins were extracted through cell scraping and sonication, and lysates were kept in 500µL lysis buffer (50mM Tris-HCl (pH7.4), 50mM NaCl, 10% glycerol, 1mM EDTA, 1mM EGTA, 5mM sodium pyrophosphate, 50mM NaF, 1% Triton X-100, cOmplete mini protease inhibitor (Roche)). The protein concentration was quantified using the Pierce BCA Protein Assay Kits (Thermo). The proteins in the samples were precipitated with 5× volume of acetone at - 20°C overnight. After washing with 1 mL acetone, the pellets were resuspended in 100µL 8M urea in 50mM TEAB buffer (pH8.0) followed by another round of quantification. The resuspended protein lysates were reduced with 10mM TCEP ((tris (2-carboxyethyl) phosphine) at 37°C for half an hour in a shaking incubator and alkylated using 55mM iodoacetamide (IAA) at room temperature in the dark. The samples were diluted to a final concentration of 1M urea using 25mM TEAB. The samples were trypsin-digested (40:1) overnight at 37°C in a shaker. To stop digestion, the samples were acidified the following day by adding pure formic acid (1% final). Digested peptides were cleaned with solid phase extraction (SPE) using Oasis HLB Cartridges (Waters),and freeze-dried overnight.

For label-free quantitative proteomic analysis, equal amounts of purified tryptic peptide mixtures were injected into a nano-UPLC system (Ultimate 3000 RSLCnano, Dionex) coupled to an Orbitrap Fusion Lumos Tribrid mass spectrometer (Thermo Fisher Scientific). Separation was performed on a 40-cm C18 column (EASY-Spray; 75µm ID, Pepsep Reprosil C18, 1.9µm particles, 120Å, 45°C). A 70-minute chromatographic gradient was applied, with mobile phase A consisting of 0.1% formic acid in water (H₂O) and mobile phase B composed of acetonitrile (ACN), water, and formic acid in an 80:20:0.1 (v/v/v) ratio. The gradient began with 2.5% mobile phase B for the first 2 minutes, then gradually increased to 35% B over 43 minutes, followed by a further increase to 45% B over the next 10 minutes. Upon reaching 45% B, the column was flushed with 99% mobile phase B for 15 minutes before returning to 2.5% B and holding for an additional 5 minutes. The injection volume for each sample was 4µL.

The mass spectrometer operated in positive ionisation mode using an EASYSpray nanosource at 2.0 kV and 275°C. Internal calibration (lock-mass) was performed in EASY-IC mode with the fluoranthene radical cation signal at m/z 202.0777. The instrument performed full MS scans (m/z 375-1500) at a resolution of 120,000 using data-dependent acquisition (DDA). In each acquisition cycle, the most intense ions (>20,000 counts) were selected for fragmentation using High-Energy Collision Dissociation (HCD, 30%). Up to 20 precursor ions were fragmented per cycle, with MS2 detection in the Orbitrap (Fusion Lumos Orbitrap; Thermo Scientific) at a resolution of 15,000. The MS2 injection time was set to 22 ms. All data were acquired using Xcalibur v4.5 software. Raw mass spectrometric data were searched using MaxQuant (v. 2.4.12.0) search engine (https://www.maxquant.org/) (Tyanova *et al*, 2016).,Tandem mass spectra were searched against the Swiss-Prot mouse (*Mus Musculus*) database. Searched data were subsequently analysed using LFQ-Analyst (https://analyst-suites.org/). Unless otherwise stated, all chemicals were from Sigma or Merck.

### cDNA constructs

The cDNA sequences for WT HINT1 for both human and mouse were codon-optimised and ordered as gene fragments (Twist Bioscience). The last 8 amino acids of the 126aa protein are: human HINT1: - RQMHWPPG*; mouse Hint1: -RQMNWPPG*. Global protein stability constructs were prepared using a pDEST GPS vector (Lin *et al*., 2018). The designed cDNA sequences were inserted by ligation into a modified pDEST4 CMV GPS vectors (Buco *et al*., 2025) and transformed in Stbl3 *E. coli*. Plasmid DNA was extracted with the ISOLATE II Plasmid Mini Kit (Bioline) following the manufacturer’s instructions. The plasmids were transfected into HEK293T cells to produce lentivirus, which was collected after 48hrs. Two cell lines were infected with the constructs – HEK293T and HEK293TK2D10, a HEK293T clone with KLHDC3 knocked out by CRISPR/Cas9 (Buco *et al*., 2025). Transduced cells were selected using puromycin and grown until they reached confluency.

### GPS reporter assay

pDEST4 HINT1 transfected HEK293T cells were assessed for dsRed and GFP expression using an inverted fluorescence microscope (Olympus IX81) using cellSens (Olympus Lifescience) software for image acquisition and/or analyzed by flow cytometry using a BD LSRIIFortessa for acquisition and FlowJo software Version 10.0 (BD Biosciences) to determine the ratio of GFP/dsRed for each reporter construct (Buco *et al*., 2025).

### Protein Extraction and Western blot analysis

For protein lysates to be used for Western blot, cultured cells were trypsinised, washed and resuspended in 150µL of RIPA buffer containing protease and phosphatase inhibitors. The samples were sonicated at 4°C in a Bioruptor Sonicating System (Diagenode) for 5 minutes, with 30-second ON/OFF pulses. Clarified samples were quantified using the Pierce BCA protein assay kit. For each sample 25µg of total protein was separated on a 4-12% Bolt Bis-Tris Plus Mini Protein Gel (Thermo Scientific) in 1X MOPS SDS Running Buffer. The proteins were transferred onto Immobilon-P PVDF and the membrane blocked with 5% Non-Fat Dry Milk (NFDM) powder in TBST (Tris Buffered Saline Tween-20) for 1 hour at room temperature. The membrane was then incubated overnight at 4°C in rabbit monoclonal anti-HINT1 antibody (Abcam, EPR5108, 1:2000) or mouse anti-Actin (Sigma Aldrich, A1978, 1:5000). The following day, the membrane was washed 3-5x using TBST, and incubated with HRP-conjugated secondary antibody for 1 hour at room temperature. The membrane was again rinsed for 3-5x with TBST and specific protein bands were then visualised using the ECL Prime Reagent (Amersham, Sigma Aldrich). Images were acquired and analysed on the iBright (Thermo Scientific).

### Data availability

The RNAseq dataset is deposited in GEO under accession codes GSE271979 (MEFs) and GSE271980 (myeloid cell lines).

### Statistical analysis

Graphs were plotted and analysed in GraphPad Prism. Statistical tests for scatter plots were done using Unpaired t-test, Ordinary One-way ANOVA, or Two-way ANOVA with multiple comparisons corrections as deemed appropriate. Comparison of non-linear curve fits were performed using F Test. All statistical legends follow these markings: ns p>0.05, *p≤ 0.05, **p≤ 0.01, ***p≤ 0.001, ****p≤ 0.0001.

## Acknowledgements

The authors thank J Heierhorst and Z Liang for comments and discussion; H-C S Yen (Academia Sinica Institute of Molecular Biology, Taipei) for GPS reporter plasmids; R Dickins (Australian Centre for Blood Diseases, Monash University) for sh*Tp53*.1224 expressing plasmid; Addgene for plasmid distribution as noted; St. Vincent’s Hospital Bioresource’s Centre for care of experimental animals; S Taylor, A Goradia, Z Liang and E Tonkin for technical assistance.

The authors acknowledge the facilities, and the scientific and technical assistance of the following Phenomics Australia nodes: Rodent Histopathology Service, University of Melbourne and Monash Genome Modification Platform (MGMP), Monash University. The *Klhdc3^+/-^* and *Klhdc3^fl/fl^* mice were via CRISPR genome editing by the Monash Genome Modification Platform (MGMP), Monash University as a node of Phenomics Australia.

## Author contributions

Conceptualization: PAVB, MFS, CRW

Methodology: PAVB, WJC-T, AH, AMC, MFS, CRW

Investigation: PAVB, WJC-T, AH, AMC, MFS, CRW

Visualization: PAB, AMC, AH, MFS, CRW

Funding acquisition: MFS, CRW

Supervision: AH, MFS, CRW

Writing – original draft: PAVB, CRW

Writing – review & editing: all authors

## Supplemental Information

**Dataset S1** – Histopathology reports

**Dataset S2** – RNA-seq of MEFs

**Dataset S3** – RNA-seq of mouse immortalised myeloid cells

**Dataset S4** – Mass spectrometry analysis of WT and *Klhdc3^-/-^*MEFs (as used in Dataset S2)

